# ATR inhibitors synergize with mitomycin C to enhance cytotoxicity in patient-derived non-muscle invasive bladder cancer organoids

**DOI:** 10.1101/2025.08.29.673024

**Authors:** Alba Zuidema, Levi Nijland, Mette Marit Vosjan, Kyah van Megesen, Bastiaan J. Viergever, Onno Kranenburg, Richard P. Meijer

## Abstract

**Background and Objective:** There is a high recurrence rate in non-muscle invasive bladder cancer (NMIBC) patients treated with intravesical chemotherapy or BCG, indicating the need for novel treatment options. Synergy between DNA damage response inhibitors and chemotherapy has been shown for different solid tumors. In this study, we explore whether combining chemotherapeutic agents with Ataxia telangiectasia and Rad3-related (ATR) kinase inhibitors is a suitable strategy to reduce recurrence in NMIBC.

**Methods:** NMIBC patient-derived organoids (PDOs; n=6) were exposed to mitomycin C (MMC), gemcitabine, or epirubicin for 2h, to mimic intravesical instillation, and subsequently to ATR inhibitors (berzosertib, ceralasertib, tuvusertib) for 72h. Cell viability and PDO regrowth potential was determined by microscopy and CellTiter-Glo® luminescence assays, during a period of 6 weeks post-treatment.

**Key Findings and Limitations:** PDO viability was severely impaired (1-22% viable cells at t=6 weeks) after combination treatment with MMC and ATR inhibitors. This effect was observed directly after treatment and maintained during 6 weeks after drug exposure. In contrast, PDOs treated with only chemotherapy or ATR inhibitors showed similar proliferation rates as untreated controls in week 6.

**Limitations:** PDO models were all chemotherapy naïve (either BCG or no pre-treatment). It would be interesting to confirm these findings in MMC-resistant models.

**Conclusions and Clinical Implications:** This study demonstrates synergy between different ATR inhibitors and MMC in a panel of six NMIBC PDOs. We propose that combining intravesical chemotherapy instillations with DNA damage response inhibitors should be further evaluated as a promising strategy to reduce recurrence rates for NMIBC.

**Plain English summary:** In this report we looked at new treatment options for non-muscle invasive bladder cancer. We discovered that tumor cells are killed very efficiently if chemotherapy is combined with drugs that prevent cells from repairing DNA damage.

## Introduction

Annually, over 120,000 people are diagnosed with bladder cancer (BC) in the EU, with over 40,000 patients dying of the disease each year. The incidence is projected to increase to 219,000 by 2030, partially due to the aging population^1,2^. The economic burden is significant, as 5% of the total healthcare costs in the EU is spent on BC^2^. Non-muscle invasive bladder cancer (NMIBC) makes up approximately 75% of all BC cases, with a high 5-year recurrence rate (∼70%) and progression to invasive disease in ∼20% of the cases^1,3^.

NMIBC patients are treated by removal of all visible tumors by transurethral resection of bladder tumor (TURB), followed by intravesical chemotherapy (first-line mitomycin C; MMC) or Bacillus Calmette-Guérin (BCG) immunotherapy^1^. These treatment options have remained largely unchanged since the 1970’s, while the high NMIBC recurrence rates and BCG shortages stress the need for novel treatment options.

Pharmacological inhibition of the Ataxia telangiectasia and Rad3-related protein serine/threonine kinase (ATR) to impair DNA damage repair (DDR), has emerged as a promising strategy in solid tumors, with multiple ATR inhibitors (ATRi) being tested in clinical studies^4^. ATR plays a crucial role in orchestrating the replication stress response, by regulating activation of cell-cycle checkpoints and stalling replication forks to allow DNA repair^4-6^. In genomically unstable cancer cells, ATRi results in the collapse of replications forks, excessive origin firing and cell cycle progression, leading to mitotic catastrophe and subsequent apoptosis^7^. ATRi synergize with DNA-damaging agents to enhance cytotoxicity ^4,7^, a rationale that has been demonstrated in pre-clinical studies of advanced BC ^8-12^. Whether ATRi in combination with MMC is a suitable strategy for NMIBC has not yet been explored.

In this study, we use patient-derived BC organoids (PDOs)^13,14^ to test the efficacy of novel ATRi combination treatments for NMIBC. Previously, we have established methods to culture BC PDOs to establish a “living biobank”^15,16^. Now, we demonstrate synergy between MMC and ATRi (berzosertib, ceralasertib, tuvusertib) in patient-derived, pre-clinical NMIBC models.

## Material and methods

### PDO culture

Ethical approval for PDO culture was granted by the Biobank Research Ethics Committee (TCBio) of the University Medical Center Utrecht (UMCU). After obtaining patient’s informed consent, organoids were isolated from patient material (either tumor tissue or urine)and cultured as previously described^15,16^. Briefly, PDOs were plated in Cultrex Reduced Growth Factor Basement Membrane Extract, Type 2, Pathclear (BME; R&D Systems Europe #3533-010-02) in pre-heated plates and maintained at 37°C in a humidified, 5% CO2 atmosphere in full culture medium (BM2+) consisting of Advanced DMEM/F-12 (ThermoFisher #12634028), supplemented with 400 µM Glutamax (Life Technologies #35050038), 10 mM HEPES (Lonza #BE17-737E), 50 U/mL–50 µg/mL penicillin–streptomycin (5.000 U/mL and 5.000 µg/mL; Life Technologies Corporation #15070063), 2% B27 supplement (Fisher Scientific #11530536), 10 mM Nicotinamide (Merck #N0636), 100 ng/ml Noggin (in-house production), 1.25 mM N-acetylcysteine (Merck #A9165), 5 μM A83-01 (MCE #HY-10432/CS-1437), 12.5 ng/mL FGF2 (Peprotech #100-18B), 25 ng/mL FGF7 (Peprotech #100-19), 100 ng/mL FGF10 (Peprotech #100-26), and 10 μM ROCK inhibitor Y-27632 (MedChem Express #HY-10583). In addition to previous protocol^16^, 0.2% primocin (Invivogen #ant-pm-05) was added to the BM2+ culture medium. PDOs were passaged in a 1:2-1:3 ratio every 1-2 weeks using TrypLE (Fisher Scientific #12604021) with a 2-min incubation at 37°C.

PDO cultures were routinely tested for mycoplasma infections using a PCR Mycoplasma Detection Set (TaKaRa #6601).

### Antibodies

Primary antibodies used are listed in Table 1. Secondary antibodies were as follows: goat anti-mouse Alexa Fluor 488 and goat anti-rabbit Alexa Fluor 647 (Invitrogen; diluted 1:200 for IF), goat anti-mouse and goat anti-rabbit HRP-conjugates (DAKO; diluted 1:2000 and 1:1000, respectively, for WB).

**Table 1.** Primary antibody list.

| Antibody | Clone | Obtained from | Host | Application |
| --- | --- | --- | --- | --- |
| KRT5 | EP1601Y | Abcam (#ab52635) | Rabbit | IF: 1:200 |
| KRT20 | Ks20.8 | DAKO (#M7019) | Mouse | IF: 1:200 |
| GATA-3 | D13C9 | Cell signaling (#5852) | Rabbit | IF: 1:200 |
| Uroplakin III | SFI-1 | Abcam (#ab78196) | Mouse | IF: 1:100 |
| phospho-Histone H2A.X (Ser139) | JBW301 | Milipore (#05-636) | Rabbit | IF: 1:400 |
| Chk1 | 2G1D5 | Cell Signaling (#2360) | Mouse | WB: 1:1000 |
| phospho-Chk (Ser345) | 133D3 | Cell Signaling (#2348) | Rabbit | WB: 1:1000 |
| GAPDH | D16H11 | Cell Signaling (#5174) | Rabbit | WB: 1:1000 |

### Confocal microscopy

PDOs (in 100 μl BME) were harvested and fixed with 4% paraformaldehyde for 10 min, washed in PBS, and permeabilized and blocked in PBS containing 1% Triton-X-100 and 5% BSA (Sigma) for 30 min. Next, PDOs were incubated with primary antibodies (Table 1) diluted in 2% BSA in PBS, overnight at 4°C. PDOs were washed three times in PBS before incubation with the secondary antibodies for 1 h at room temperature. After three washing steps, slides were mounted using ProLong™ Gold Antifade Mountant with DNA Stain DAPI (Invitrogen # P36931). Images were obtained at room temperature using a Zeiss LSM700 confocal microscope, controlled with ZEN 2011 software (Version 14.0.21.201), with a 40x (NA 1.2) water or 63x (NA 1.4) oil objective.

### Western blotting

Three-day old PDOs (in 200 μl BME/condition) were treated for 2h with 2-4 μM MMC, following 3 wash steps in PBS and 20h incubation in BM2+. Next, PDOs were treated for 3 days with berzosertib (TargetMol #T2669) in BM2+. PDOs were washed 3x in PBS, incubated with 1mg/ml Dispase (Gibco #17105041) for 30 min at room temperature, and harvested in eppendorfs. Cells were lysed in Laemmli lysis buffer (12% 1M Tris pH 6,8, 20% 95%-Glycerol, 12,5% 20%-SDS in H20) and boiled (100°C) for 10 min. Protein concentrations were determined using a Lowry assay. Per lane, 15-20 µg of protein was loaded on pre-cast 1,5mm 4-12% NUPAGE Bis-Tris mini protein Gel (Invitrogen #NP0336BOX). Protein was transferred to PVDF transfer membranes (Bio-Rad #1704156) using the Trans-Blot® Turbo™ Transfer System (Bio-Rad #1704150). Membranes were blocked for 1h in 5% non-fat dry milk in TBST at room temperature. Primary antibodies were dissolved in 1% non-fat dry milk in TBST and incubated overnight at 4°C. Membranes were washed 3 x 10 min in TBST and incubated with secondary antibodies for 1h at room temperature. After washing 3 x 10 min in TBST and 10 min in TBS, protein expression was visualized using ECL Western Blotting detection kits (Cytiva #RPN2209 and #RPN2232), on a Amersham Imager 600 (Cytiva).

### Drug screens

PDOs were harvested from BME droplets, dissociated using TrypLE (3 min incubation at 37°C), and filtered using a 70 µm-cell strainer to obtain a single cell suspension. In a 96-well plate, 3000 single cells per well were plated in 4 µl BME droplets and maintained in BM2+ culture medium. After 3 days, PDOs were treated with MMC for 2h, as indicated in figures. Next, PDOs were washed three times with PBS, before BM2+ culture medium was added, and 20h later berzosertib, ceralasertib (TargetMol #T3338), or tuvusertib (TargetMol #T10406) was added for 3 days. Next, PDOs were washed in PBS (3x) and maintained in BM2+, which was refreshed twice per week. Images were taken weekly using a CellInsightTM CX5 High Content Screening Platform (Thermo Scientific) with Thermo Scientific HCS Studio software (version 6.6.1). Cell viability was measured using CellTiter-Glo 3D (Promega #G9683) assay, according to manufacturers’ instructions, and read out on a GloMax Navigator (Promega) or Spectramax i3 (Molecular devices) plate reader.

### Image quantifications

PDO images were segmented using Ilastik (version 1.4.0) Pixel Classification Workflow^17^, using 5 representative images per organoid line for training. Segmented images were further analyzed using Fiji (ImageJ) to measure total organoid area per image^18,19^.

### Statistical analysis

Kruskal–Wallis with Dunn’s multiple comparisons test was performed using GraphPad Prism (version 10.5.0). In figures, statistically significant values are shown as ^*^P < 0.05; ^**^P < 0.01; ^***^P < 0.001; ^****^P < 0.0001.

## Results

### NMIBC PDO panel

PDO models were selected from a previously established “living biobank” at the University Medical Center Utrecht, which contains BC organoids derived from urine samples or bladder tumor tissue biopsies (TURB or cystectomy)^16^. These PDOs resemble the histopathological and genetic features of the original tumor tissue and retain a comparable level of heterogeneity after isolation^16^.

The six PDO models were derived from chemotherapy-naïve NMIBC patients with tumors that displayed diverse pathological features, including carcinoma in situ (CIS), low- or high-grade Ta/T1 tumors (Table 2; Fig.1). Morphologically, most PDOs displayed a round, solid phenotype, without a central cavity. Organoids containing a luminal space could occasionally be observed, in particular in the UBTOR36.3 line. The UBTOR8 showed a dense, grape-like morphology, with partial growth in 2D. Urothelial carcinoma (UC) origin of the PDOs was confirmed by visualizing the expression of uroplakin III and GATA3 using immunofluorescence staining (Fig.1). Most PDOs showed mixed expression of basal (keratin 5) and luminal (keratin 20) markers, indicating that the intra-tumoral heterogeneity observed in the original tumor was maintained during long-term culture (Fig.1)^16^. On a cellular level, either KRT5 or KRT20 was expressed (Fig.1).

**Figure 1.**
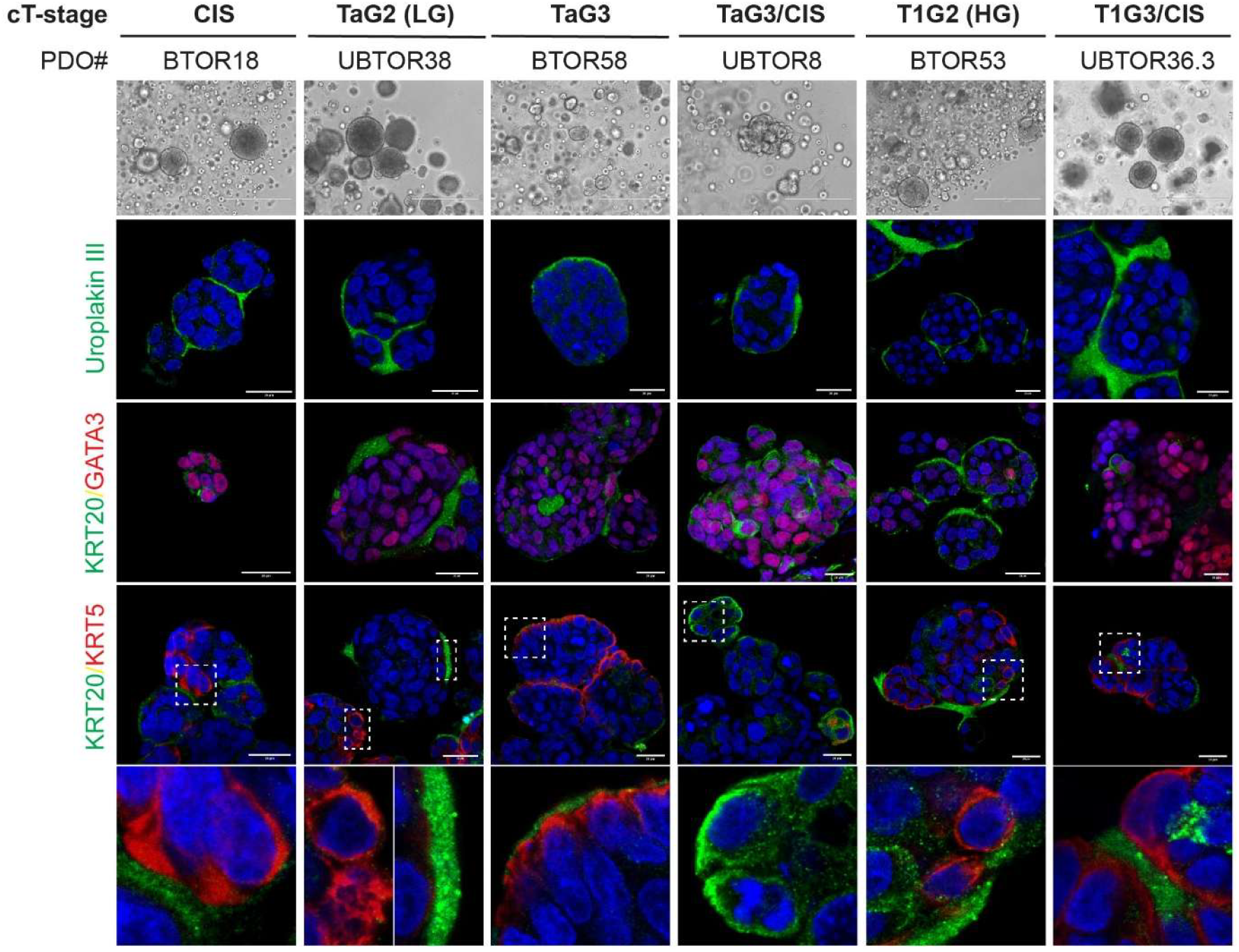
Expression of urothelial carcinoma markers in NMIBC organoids. Representative brightfield (top panel) and immunofluorescence images showing uroplakin III (green) and keratin 20 (KRT20; green) with GATA3 (red) or keratin 5 (KRT5; red) in NMIBC patient-derived organoids (PDOs). Nuclei were stained with DAPI (blue). Scale bars: 400 μm (brightfield) and 20 μm (IF). **BTOR** = tissue-based bladder cancer organoid line; **UBTOR** = urine-based bladder cancer organoid line; **CIS** = carcinoma in situ; **LG** = low grade; **HG** = high grade.

**Table 2.** Patient information NMIBC PDO panel.

|  | Patient information |  |  |  |  |  |
| --- | --- | --- | --- | --- | --- | --- |
| PDO# | Gender | Age | Classification | Grade | cT-stage | Pre-treatment |
| BTOR18 | F | 72 | UC | G3 (HG) | CIS | BCG |
| UBTOR38 | M | 70 | UC | G2 (LG) | TaG1-2 | - |
| BTOR58 | M | 86 | UC | G3 (HG) | TaG3 | - |
| UBTOR8 | M | 58 | UC | G3 (HG) | TaG3/CIS | BCG |
| BTOR53 | F | 64 | UC | G2 (HG) | T1G2 | BCG |
| UBTOR36.3 | M | 62 | UC | G3 (HG) | T1G3/CIS | BCG |
| **BTOR = tissue-based bladder cancer organoid line; UBTOR = urine-based bladder cancer organoid line; CIS = carcinoma in situ** |  |  |  |  |  |  |

### Berzosertib inhibits MMC-induced ATR signaling

NMIBC PDOs (Table 2; Fig.1) were used to evaluate the efficacy of ATRi combined with chemotherapy. First, PDOs were exposed to MMC-treatment for 2h, in order to mimic intravesical treatment. This treatment induced DNA damage in the majority of tumor cells, as indicated by the presence of γH2AX (Fig.2A,B)^20^. To explore whether combination treatment with ATRi would result in synergistic antitumor activity, MMC treatment was followed by administration of berzosertib (formerly M6620, VX-97)^21^. Using western blot, activation of ATR signaling after MMC treatment was demonstrated, which resulted in the phosphorylation of the downstream target Chk1 at Ser-345 (Fig.2C,D). Berzosertib efficiently blocked ATR signaling when NMIBC PDOs were exposed to the drug for 3 days, with the treatment starting 20h after MMC-treatment (Fig.2C,D).

**Figure 2.**
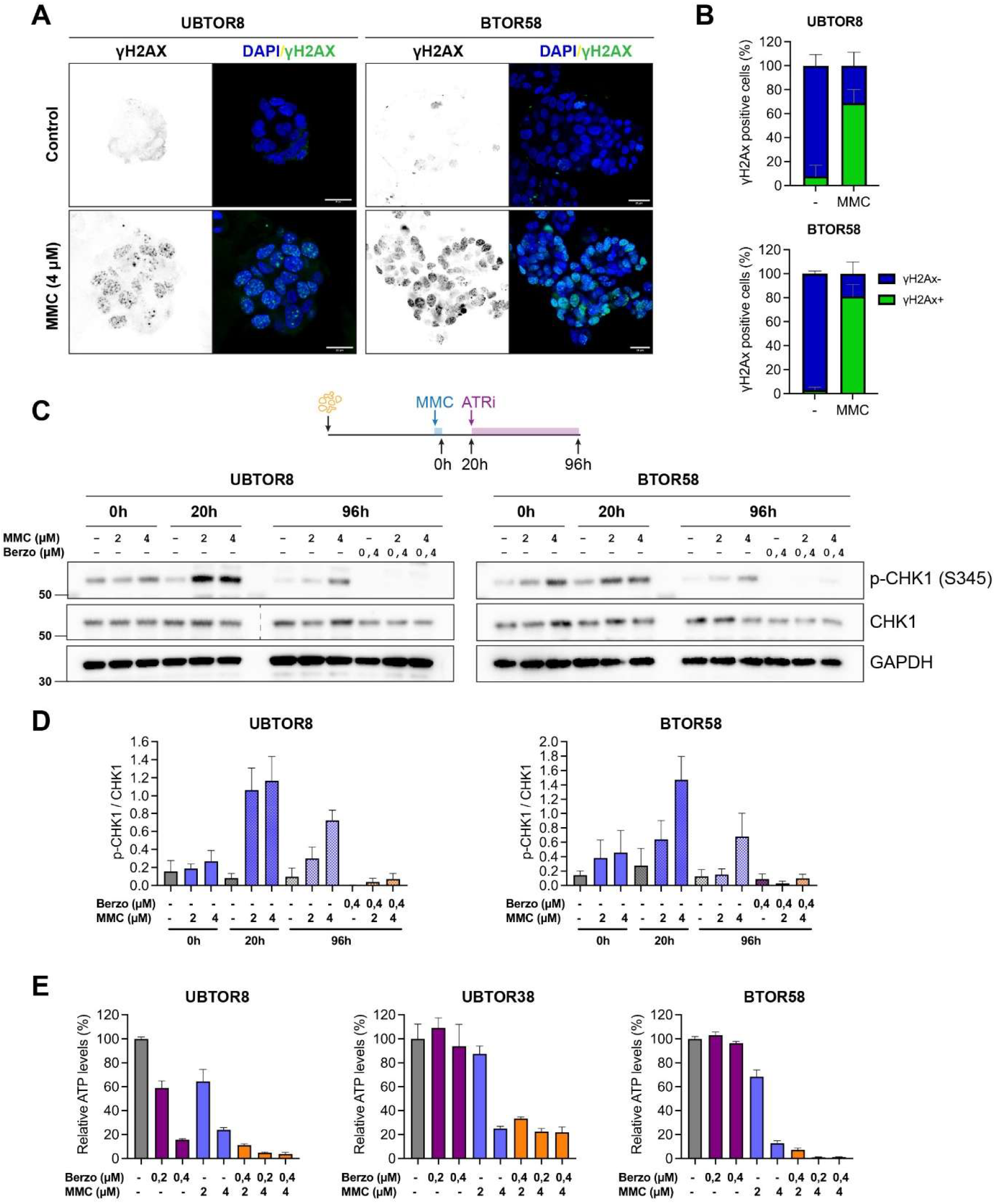
Berzosertib inhibits MMC-induced ATR signaling. (**A**) Immunofluorescence images showing nuclei (DAPI staining; blue) with yH2AX foci (green in merged images) in 4 μM MMC-treated and untreated controls. PDOs were treated with MMC for 2h and fixed 20h later in paraformaldehyde. Scale bars: 20 μm (**B**) Quantifications of the percentage of yH2AX positive cells per image (based on 6 different fields per PDO per condition). (**C**) PDOs were treated with MMC (0, 2 or 4 μM, as indicated) for 2h and 20h later with 0,4 μM berzosertib for 72h. Cell lysates were made directly after MMC-treatment (t=0) and before (t=20h) and after berzosertib addition (t=96h). Representative western blots of phosphorylated Chk1 (Ser345), total Chk1, and GAPDH (loading control) are shown, with quantification of signal intensities (mean with SD) of 2 independent experiments per PDO shown in (**D**). (**E**) Cell viability measurements (Cell Titer Glo) directly after (combination) treatment at t=96h. Bar graphs show mean values with SEM normalized to untreated controls (n=4 technical replicates per condition).

Cell viability measurements conducted directly after treatment confirmed that the MMC-berzosertib combination is more effective at eradicating tumor cells than MMC monotherapy (Fig.2E). In particular, synergy could be observed for UBTOR38 organoids treated with 2 µM MMC and 0,4 µM berzosertib (Relative ATP levels >88% for mono treatment versus 33% for combination) and for BTOR58 (Relative ATP levels >68% mono treatment versus 7% for combination). At the highest drug concentrations used, the combination treatment resulted in almost complete cell killing (<5% viable cells) in UBTOR8 and BTOR58 organoids. The untreated UBTOR38 organoids grew slower in culture, which could explain why the cell viability was ∼30% after treatment.

Taken together, berzosertib inhibits ATR-mediated DDR, which promotes cell death in NMIBC PDOs after MMC treatment.

### PDO viability is long-term impaired after MMC-berzosertib combination treatment

Next, we evaluated whether ATRi could prevent regrowth of chemotherapy-treated PDOs. To this end, we exposed six PDOs to a 2-hour MMC-treatment and, subsequently, to berzosertib for 72h (as in Fig.2C-E). Regrowth potential was assessed by microscopy during a 6-week period after mono (MMC or berzosertib) or combination treatment and compared to the growth of untreated PDOs. Without exception, combination treatment impaired the growth of NMIBC PDOs and prevented regrowth (Fig.3). This antitumor effect was maintained during the 6-week time course, while PDOs treated with only MMC or berzosertib showed comparable growth (after an initial growth delay) as the untreated controls in week 6 (Fig.3). These results were confirmed by measuring the amount of metabolically active cells, based on ATP-levels, at week 6 post-treatment (Fig.3).

**Figure 3.**
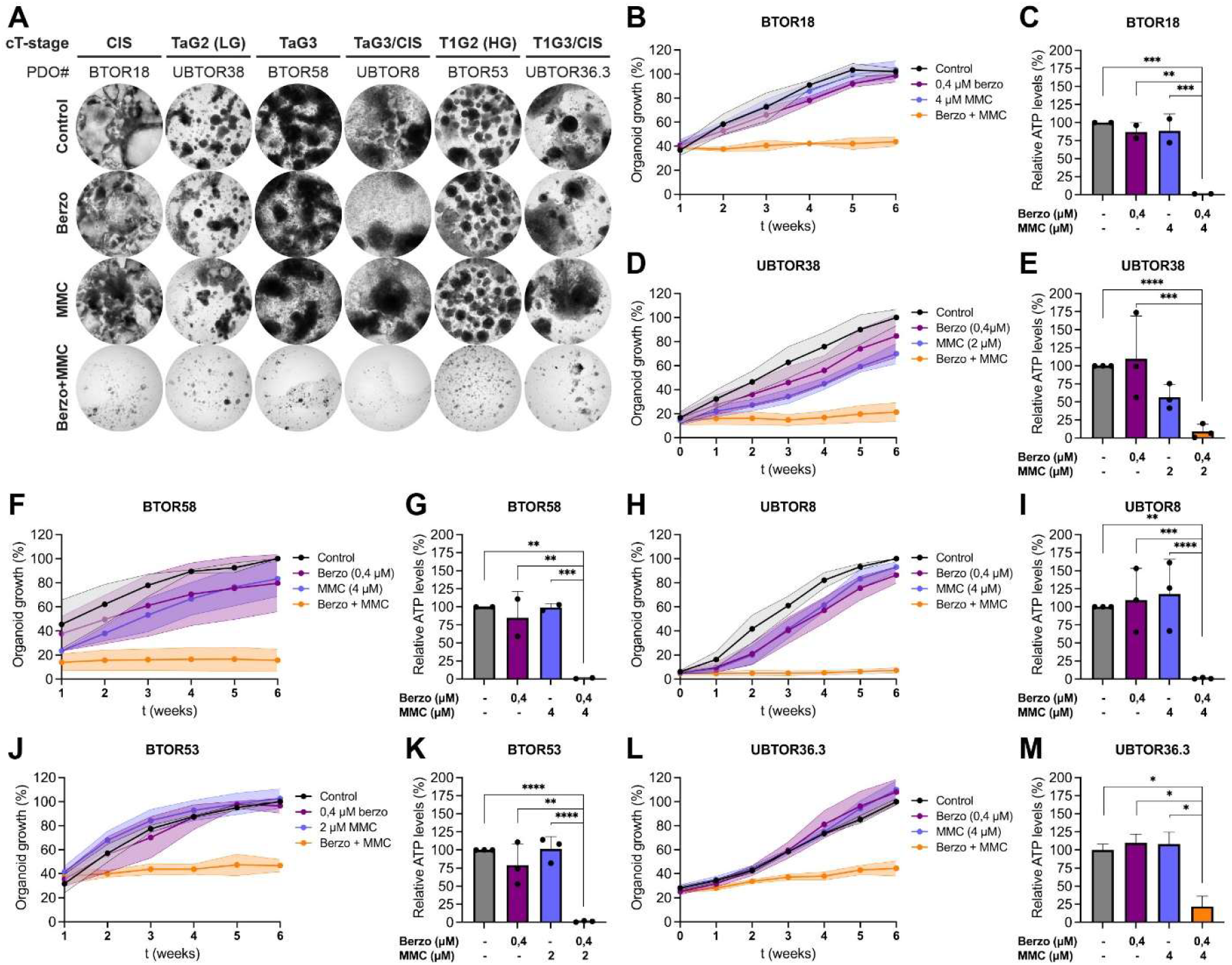
Berzosertib synergizes with MMC to prevent regrowth of NMIBC PDOs. (**A**) PDOs were treated with 2-4 μM MMC and/or 0,4 μM berzosertib, following the timeline described in Fig.2C. Weekly images were taken and representative images at t=6 weeks after treatment are shown for each condition. (**B**,**D**,**F**,**H**,**J**,**L**) Image quantifications of total PDO area per image, treated as indicated, and normalized to untreated controls at week 6 (=100%). (**C**,**E**,**G**,**I**,**K**,**M**) Cell viability measurements (Cell Titer Glo) at t=6 weeks. Bar graphs show mean values with SD normalized to untreated controls. Dots in the bar graphs represent the mean value per biological replicate (n=2 for BTOR18, BTOR58; n=3 for UBTOR8, UBTOR38, BTOR53) with n≥3 technical replicates per condition per independent experiment. Kruskal–Wallis with Dunn’s multiple comparisons test was performed to determine statistical significance, ^*^P < 0.05; ^**^P < 0.01; ^***^P < 0.001; ^****^P < 0.0001.

In these experiments, PDOs were exposed to MMC at concentrations ranging from 2-20 µM (Fig.3; S1). Synergy between ATRi and MMC could be best observed at MMC concentrations of 2 µM (UBTOR38, BTOR53; Fig.3D-E,J-K) and 4 µM (UBTOR8, BTOR18, UBTOR36.3, BTOR58; Fig.3B-C,F-I,L-M), at which DNA damage was present in the majority of tumor cells and ATR signaling was activated (Fig.2A-D). To achieve the same growth inhibition with MMC monotherapy at week 6 post-treatment, a 4-5 fold higher MMC concentration was needed (Fig.S1).

**Figure 4.**
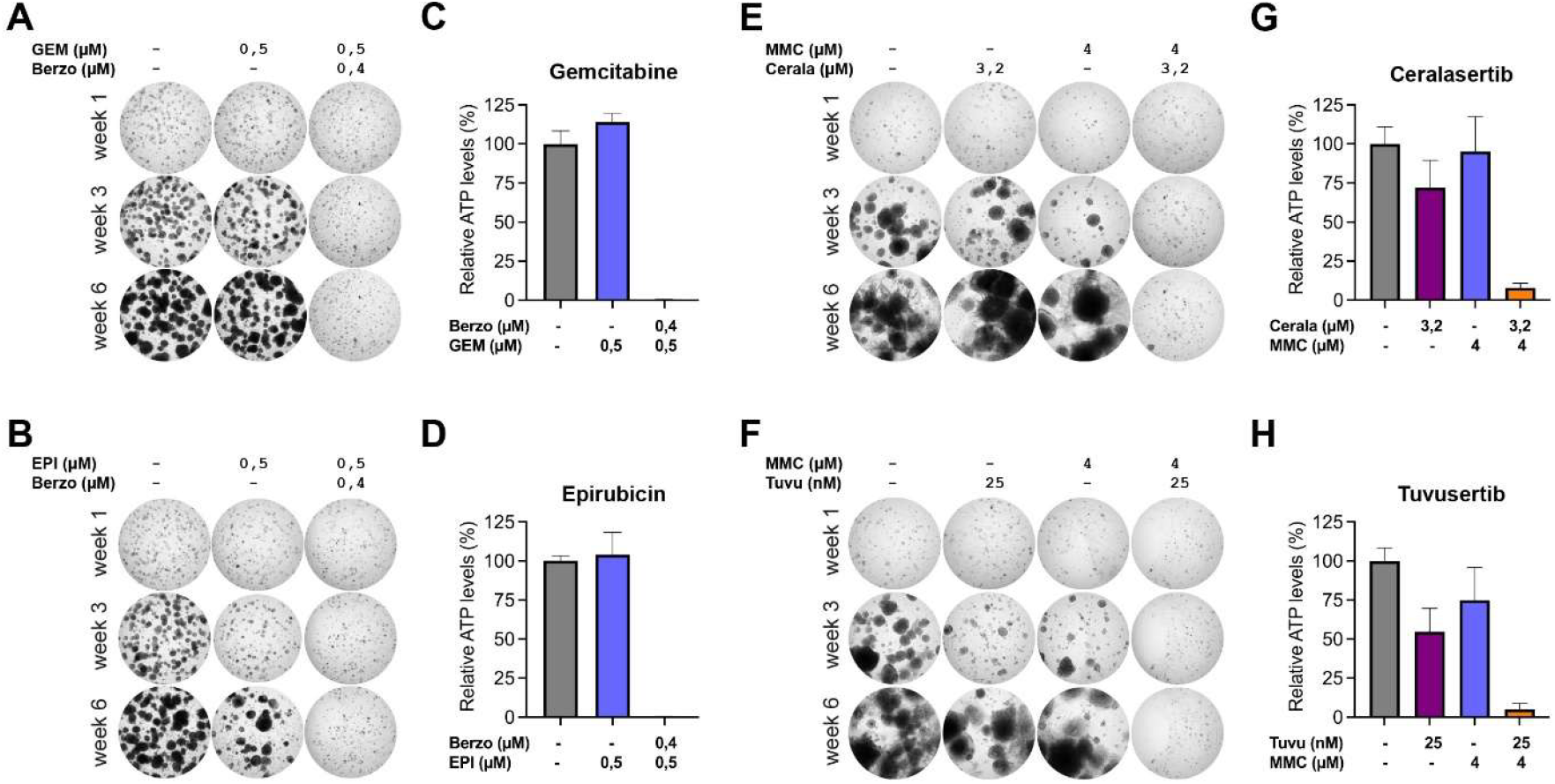
ATR inhibition following chemotherapy eradicates NMIBC cells. (**A**,**B**) BTOR58 organoids were treated for 2h with 0,5 μM gemcitabine (GEM) (**A**) or 0,5 μM epirubicin (EPI) (**B**) alone or in combination with 0,4 μM berzosertib follow-up treatment for 72h. Weekly images were taken for 6 weeks after treatment and representative images at t=1, 3, and 6 weeks are shown. Cell viability assays at t=6 weeks are shown in (**C**,**D**). (**E**,**F**) UBTOR8 organoids were treated for 2h with 4 MMC, alone or in combination with 3,2 μM ceralasertib (**E**) or 25 nM tuvusertib (**F**) follow-up treatment for 72h. Representative images at t=1, 3, and 6 weeks after treatment are presented. Cell viability assays at t=6 weeks are shown in (**G**,**H**). Bar graphs represent mean with SD (n=3).

Taken together, we demonstrated synergy between berzosertib and MMC in a panel of NMIBC organoids.

### ATRi prevents regrowth of NMIBC PDOs treated with DNA damaging agents

To analyze if the observed synergy between berzosertib and MMC would be applicable to other DNA-damaging agents, BTOR58 organoids were exposed to gemcitabine and epirubicin mono- or combination treatment with berzosertib (Fig.4A-D). In line with a previous study^9^, berzosertib severely impaired cell viability and regrowth potential in combination with gemcitabine and epirubicin (Fig.4A-D). Finally, we compared berzosertib to other inhibitors that are currently evaluated in clinical studies: ceralasertib and tuvusertib^22,23^. Again, we observe a synergistic effect between MMC and ATRi (Fig.4E-H).

In conclusion, synergy between ATRi and MMC can be exploited to eradicate NMIBC cells and prevent regrowth.

## Discussion

In this study, we demonstrate that ATR inhibitors synergize with MMC to enhance its cytotoxicity in NMIBC PDOs. The rationale of combining DNA damage response inhibitors (DDRi; e.g. ATM, ATR, CHK1, CHK2, DNA-PK, PARP, WEE1) and chemotherapy has been explored for different solid tumors with encouraging outcomes^24-27^. Based on our promising results in patient-derived models, we propose that combining intravesical chemotherapy instillations with these inhibitors should be further investigated as a strategy to reduce recurrence rates for NMIBC.

Lack of patient-representative models for novel drug screening and identification of targeted therapies has delayed preclinical NMIBC research. The use of PDOs is finally gaining traction in BC research, with several studies demonstrating that these models recapitulate histopathological features and genomic profiles of the bladder tumor, while retaining a comparable level of heterogeneity after isolation^13,15,16,28-33^. Most BC organoid research has focused on advanced UC. In addition, we built a NMIBC biobank that includes preclinical models for CIS, as treatment options for these patients are limited only to BCG after TURB^1^. Our findings that ATRi strongly promotes MMC cytotoxicity also in CIS-derived organoids (UBTOR8, BTOR18, UBTOR36.3), might open new therapeutic avenues for these patients.

DDRi is a very active area of research, with various inhibitors in clinical studies for multiple tumor types. However, only one randomized phase 2 trial (Clinical-Trials.gov identifier: NCT02567409) has explored the benefit of combined cisplatin and gemcitabine plus berzosertib versus cisplatin and gemcitabine alone in patients with advanced UC. Unfortunately, the findings of this trial indicated that berzosertib had no added benefit compared to cisplatin and gemcitabine chemotherapy, likely because of adverse hematologic effects resulted in substantial dose reductions during the trial^34^. Although berzosertib did show promising results in combination with gemcitabine in platinum-resistant ovarian cancer^35^, its clinical development has been discontinued in favor of the oral ATR inhibitor tuvusertib (M1774)^22^. Tuvusertib suppressed cancer cell viability at nanomolar concentrations in vitro, demonstrating enhanced potency compared to berzosertib and ceralasertib, while showing a manageable safety profile in the DDRiver Solid Tumors 301 phase I study^22,36^. Our findings confirm that tuvusertib is more potent than berzosertib and ceralasertib in eradicating NMIBC cells (Fig.4). Other ongoing trials evaluating ATRi in combination with chemotherapy include: ceralasertib (NCT03669601), ART0380 (NCT04657068), camonsertib (NCT04497116), and elimusertib (NCT04616534).

Finally, we propose that for the treatment of NMIBC, ATRi could be administered intravesically, e.g. using a sustained drug-delivery system like TAR-200^37^, to overcome adverse effects observed during systemic ATRi treatment in combination with chemotherapy.

Upon inhibition of ATR, cells might use alternative pathways to repair DNA. If these other DDR pathways are simultaneously inhibited as ATRi (either intrinsically by loss-of-function mutations or pharmacologically), synthetic lethality takes place. This effect has been described, for example, for BRCA1/2-defective cells treated with PARPi and for p53-deficient cells with ATMi/PARPi^27^. Moreover, mutations in ARID1A, which are common in BC, lead to synergy with ATRi^38,39^. A recent study showed that ARID1A-deficient BC PDOs can be selectively eliminated by the Chk1 inhibitor prexasertib (LY2606368), indicating the potential of PDOs for biomarker research and personalized medicine^40^.

## Conclusions

This study demonstrates synergy between ATRi and MMC in NMIBC PDOs. Therefore, it could be a promising strategy to combine intravesical chemotherapy instillations with DDRi for NMIBC patients.

## Acknowledgements

We acknowledge the Cell Microscopy Core (CMC) of the Center for Molecular Medicine, UMC Utrecht for providing microscopy training and service.

## Author contributions

Richard P. Meijer had full access to all the data in the study and takes responsibility for the integrity of the data and the accuracy of the data analysis.

*Study concept and design*: Zuidema, Nijland, Van Megesen, Kranenburg, Meijer.

*Acquisition of data*: Zuidema, Nijland, Vosjan.

*Analysis and interpretation of data*: Zuidema, Nijland, Vosjan, Kranenburg, Meijer.

*Drafting of the manuscript*: Zuidema, Meijer.

*Critical revision of the manuscript for important intellectual content*: Kranenburg, Meijer.

*Statistical analysis*: Zuidema.

*Obtaining funding*: Meijer.

*Administrative, technical, or material support*: Viergever

*Supervision*: Zuidema, Kranenburg, Meijer.

*Other*: None.

## Supplementary Figure

**Figure S1.**
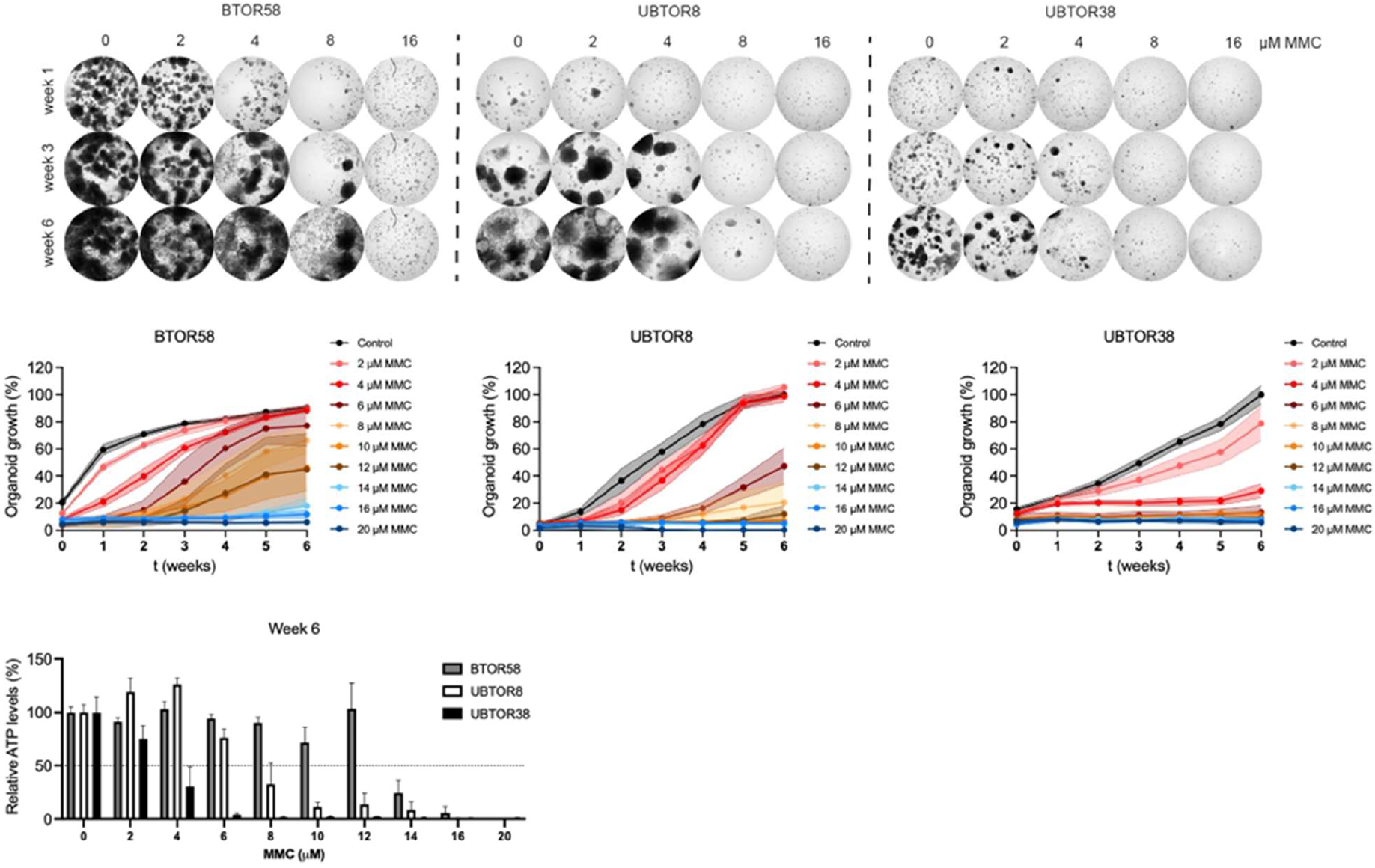
NMIBC PDO treated with MMC. PDOs were treated for 2h with MMC as indicated. Weekly images were taken for 6 weeks after treatment and representative images at t=1, 3, and 6 weeks are shown. Cell viability was measured at t=6 weeks. Bar graphs represent mean with SD (n=3).

